# Cellular heterogeneity results from indirect effects under metabolic tradeoffs

**DOI:** 10.1101/558619

**Authors:** Aina Ollé-Vila, Ricard Solé

## Abstract

The emergence of multicellularity requires the coexistence of diverse cellular populations displaying cooperative relationships. This enables long-term persistence of cellular consortia, particularly under environmental constraints that challenge cell survival. Toxic environments are known to trigger the formation of multicellular consortia capable of dealing with waste while promoting cell diversity as a way to overcome selection barriers. In this context, recent theoretical studies suggest that an environment containing both resources and toxic waste can promote the emergence of complex, spatially distributed *proto-organisms* exhibiting division of labor and higher-scale features beyond the cell-cell pairwise interactions. Some previous non-spatial models suggest that the presence of a growth inhibitor can trigger the coexistence of competitive species in an antibiotic-resistance context. In this paper we further explore this idea using both mathematical and computational models taking the most fundamental features of the *proto-organisms* interactions. It is shown that this resource-waste environmental context, in which both species are lethally affected by the toxic waste and metabolic tradeoffs are present, favours the maintenance of diverse populations. A spatial, stochastic extension confirms our basic results. The evolutionary and ecological implications of these results are outlined.

## I. INTRODUCTION

Multicellularity has evolved independently at least 25 times in our planet[1–3]. The rise of multicellular systems provided the fabric for further evolutionary events and additional layers of complexity. It became a crucial step towards the rise of complex ecosystems and provided novel opportunities for new niches. It was also a first step towards complex organisms with diverse tissues and organs as well as sensors and actuators. In a different (but deeply related) context, most biotransformations that occur in nature are carried out by microbial consortia with different species playing different tasks and often associated in some kind of spatial arrangement (Fig. 1a). In both cases, multicellular assemblies arise as a novel way to deal with environmental sources of stress.

**FIG. 1:**
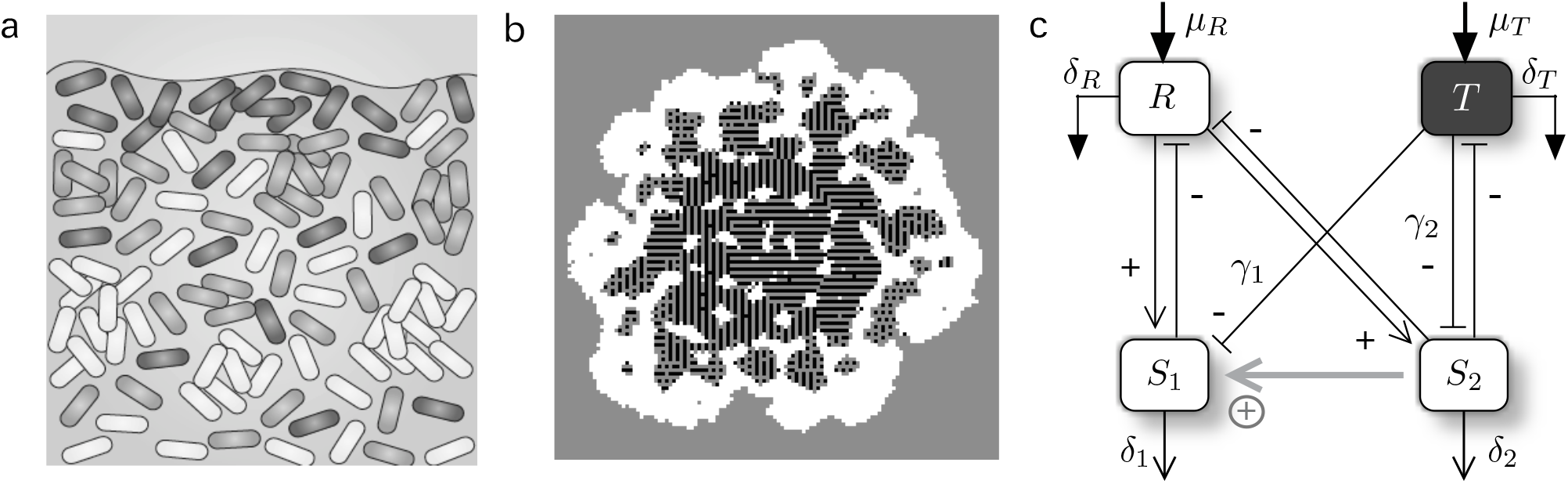
Multicellular structures in a resource-waste environment. In many different scenarios, microbial biofilms (a) are formed as a way to protect bacterial communities from selective barriers. They typically involve heterogeneous consortia with division of labour. In a different, butt related context, complex proto-organismal structures have been obtained (b) in an evolutionary spatial model of cell populations with two basic types (white and black squares, respectively) in a medium (gray sites) including both diffusing nutrients and toxins. The cells evolve specific adhesion patterns that sustain complex, repeatable structures (see Duran-Nebreda et al.[12] for details). A mean field model (c) aims at capturing the most fundamental features of the PRO model in order to understand the origins of such complex structures. The non-direct interaction (gray bottom arrow) builds up a positive link which has a critical role in maintaining cellular heterogeneity.

One particular instance that has been widespread is the need of dealing with toxic waste, as it can deeply affect the growth potential of cell populations and even determine their decline and extinction. As a result, specialised mechanisms of degradation or neutralisation have evolved multiple times (refs). Not surprisingly, important evolutionary transitions in our past Biosphere involved dealing with threshold concentrations of toxic or damaging chemicals or radiation levels[4]. Thriving in such selective environments, whether associated physical factors, substrate heterogeneity, shortening of essential toxins or antibiotics, pervades multiple behavioural and self-organising patterns that require the maintenance of a stable multicellular consortium[5–9]. Likewise, certain tradeoffs are also important evolutionary drivers, as it has been shown for the trade-off between rate and yield of ATP production[10, 11]. In this case, opposite strategies are selected depending on the environmental context, with cooperation and competition having a critical role in the evolutionary outcome.

Models for the origins of MC require a minimal set of assumptions capable of capturing the essential mechanisms required for the maintenance of cell diversity. Such coexistence should be a consequence of cooperative interactions enhancing diversity while providing resilience in front of environmental fluctuations. In a recent paper[12], it was shown that complex proto-organismal (PRO) structures can emerge in a resource-waste environment, as shown in Fig. 1b. In this spatial model, two basic cell types were introduced, linked through a stochastic switching mechanism. One type was able to use resources and replicate, whereas the other was able to process waste at the expense of a reduced replicative potential. Both cell types where affected by the toxic waste through direct cell mortality. The model also incorporated evolutionary dynamics and cell-cell adhesion mediated by an energy minimisation rule that plays a relevant role in a broad range of morphogenetic phenomena [13] and references therein. Despite the lack of a gene regulatory network supporting a true developmental program, the model was able to generate complex, spatially self-organised patterns far beyond a simple clustered, heterogeneous mixture of cells.

What is the fundamental origin of these proto-organisms? What pervades the strong interdependence between cell types? Previous studies on the dynamics of bacterial populations in a chemostat under the presence of antibiotics[14–16] showed that stable coexistence can be achieved under the presence of external inhibitors (mediated by plasmids). Similarly, studies by Eshel Ben-Jacob and co-workers has shown that complex cooperative behavioral patterns (including spatial patterns) emerge out of cooperative responses to environmental stress [17, 18]. These studies reveal the power of self-engineering associated to antibiotic resistance[19]. Here we present a mean field model capturing the most relevant features of the spatial PRO model in order to understand the possible key features predating the observed proto-organisms. We find the necessary conditions for the heterogeneous population to exist under the studied scenario, which involve a set of asymmetries and positive correlations allowing us to provide novel insights into the field of toxic-mediated complexity

## II. RESOURCE-TOXIN-CONSUMERS MODEL

As described by the diagram of Fig. 1c, our two cell populations exploit the resource *R* as consumers while both are damaged by the toxic waste *T*. Since the original inspiration of the model[12] involves closely related cell types, several parameters are taken equal, although as a first approximation we do not link the two cell populations through any developmental nor mutual switching mechanism. However, trade-offs impose some asymmetries. Two types of cells are considered. The first (*S*_1_) replicates at a given rate under the presence of *R* and is lethally affected by the presence of *T*. On the other hand, the second (*S*_2_) can degrade the toxic component *T* while showing a reduced replication potential. Such trade-off would result from the energy investment associated to *T* degradation, which necessarily implies that the *S*_2_ species will be less efficient in replicating. Importantly, this toxic degrading species also experiences the lethal inhibition from toxic waste.

The full set of equations defining our mean field model are now:

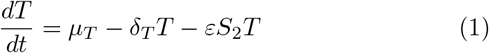

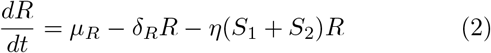

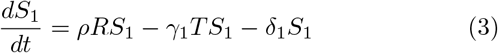

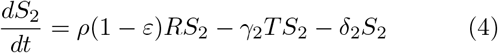

where *µ*_*T,R*_ are the influx rates for toxic *T* and resources *R*, respectively, *δ*_1,2_ are the cell mortality rates, *ρ* is the basal cell replication efficiency, *ε* is the efficiency of toxic degradation by *S*_2_, which is reflected in a lower efficiency for replication as a trade-off (*ρ*(1 − *ε*)), *η* is the rate of resource degradation by both species and *γ*_1,2_ are the mortality rates due to toxic for each species.

The previous set of equations has four relevant fixed points 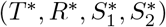, namely the full extinction point,

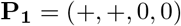

the coexistence point with both consumers present,

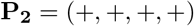

and two single-survivor fixed points, namely:,

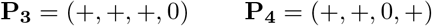

(see S-V SM for the full expressions). Below (and in S-V and S-VI SM) we study the stability of these points and determine the parametric conditions associated to each.

The model shares some commonalities with the classical one resource-two consumer (1R2C) model[20], where the Competitive Exclusion Principle[21] rules out the possibility of coexisting species. However, the presence of a toxic in the medium changes this scenario, creating an indirect positive effect from *S*_2_ to *S*_1_ (grey line in Fig. 1c), which rescues the coexistence scenario. We will show how this indirect positive link, the direct cell mortality through toxic endured by both cells and the metabolic tradeoff of species *S*_2_ impose certain constraints in the form of asymmetries and positive correlations that must hold in order to rescue the coexistence scenario.

## III. RESULTS

### A. Mean field model

The stability analysis of the subsystems *RTS*_1_ and *RTS*_2_ is properly presented in S-I, II, III SM leading to a proper intuition to further study the complete system. As the stability analysis of the mean field model is nearly intractable, in order to assess the stability of the presented system we have made use of qualitative stability analysis tools[22–26] (see S-VI SM). These allow us to show that there are some conditions which do not guarantee the stability of our particular system for any parameter values (see also S-VI SM for more details). This fact requires a deeper analysis of the system and below we present the conditions that need to hold for our system to be stable. The first is the need of an asymmetry in the toxic sensitivity values *γ*_1_*/*_2_ (see Fig. 2). The second demonstrates the critical role of two of the system links which build up an indirect positive relation: these two links need to hold a positive correlation so that coexistence is possible (see Fig. 3a,b). Moreover, our system displays a metabolic tradeoff regulated by *ε*, and this is also constraining the system’s behaviour in enabling its full viability (see Fig. 3c). These conditions are fully developed below.

**FIG. 2:**
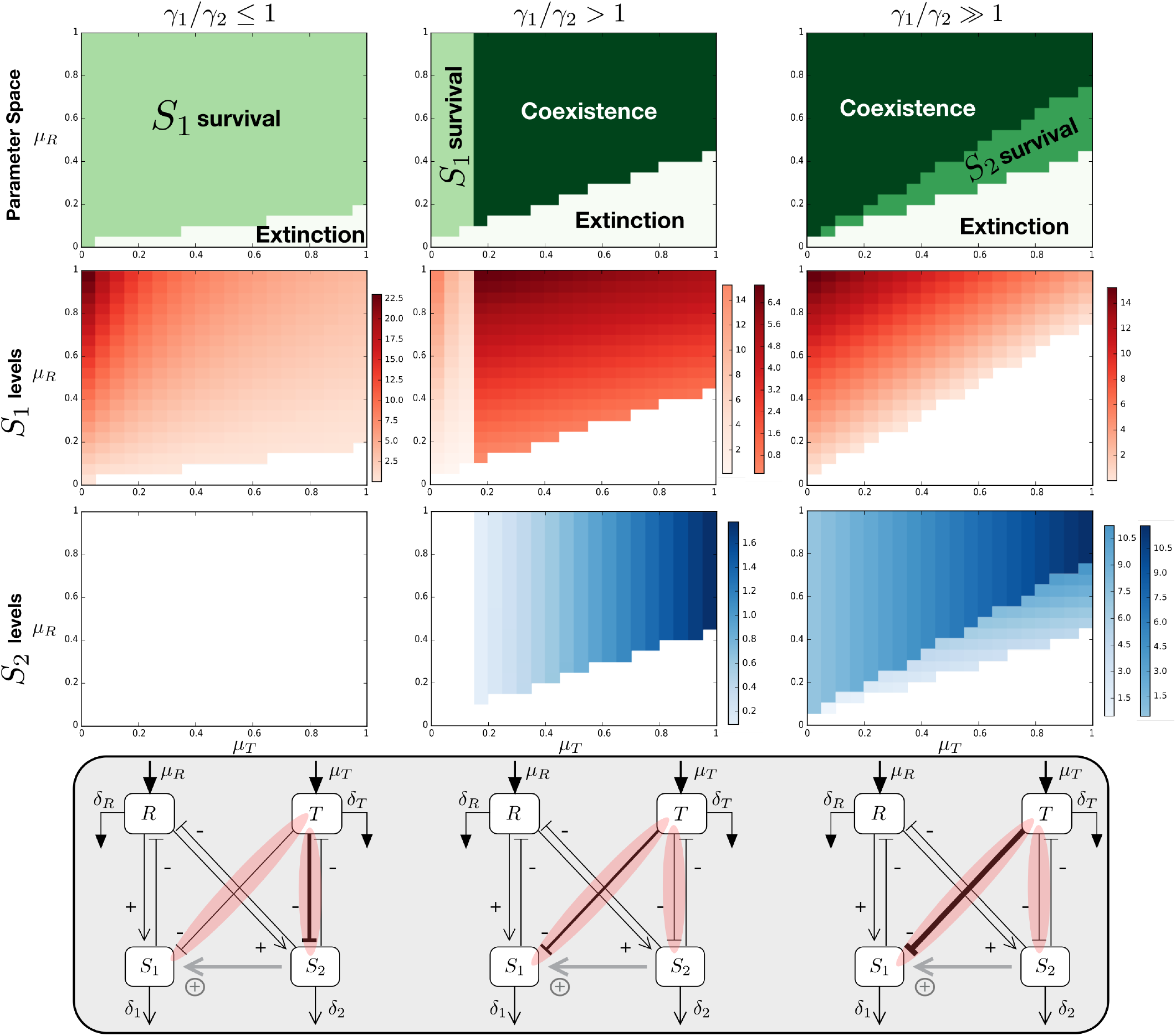
Parameter space of the systehm under different conditions of *γ*_1_*/γ*_2_ (each column) as a function of *µ*_*T*_ and *µ*_*R*_. This means that the Toxic substance needs to have a more harmful effect on species *S*_1_ in order that the presence of species *S*_2_, which is able to process this Toxic substance, increases the fitness of the whole system allowing the coexistence of the two species. We observe how the relative value of these two parameters is determining the absence or presence of coexistence and regions of *S*_1_ and *S*_2_ survival. In the second and third rows we observe the value of the fixed point of *S*_1_ and *S*_2_, respectively, using different gradients for the different regions of the parameter space shown in the first row. The values for *R* and *T* in the fixed points for these regions can be found at the Supplementary Material. For *γ*_1_*/γ*_2_ *≤*1, *γ*_1_ = *γ*_2_ = 0.1, for *γ*_1_*/γ*_2_ > 1, *γ*_1_ = 0.17 and *γ*_2_ = 0.1 and for *γ*_1_*/γ*_2_ »1, *γ*_1_ = 0.3 and *γ*_2_ = 0.1. The rest of the parameters used (all shared in the three parameter spaces) are: *ρ* = 0.3, *ε* = 0.3, *η* = 0.1, *δ*_*R*_ = *δ*_*T*_ = 0.1, *δ*_1_ = *δ*_2_ = 0.1.

**FIG. 3:**
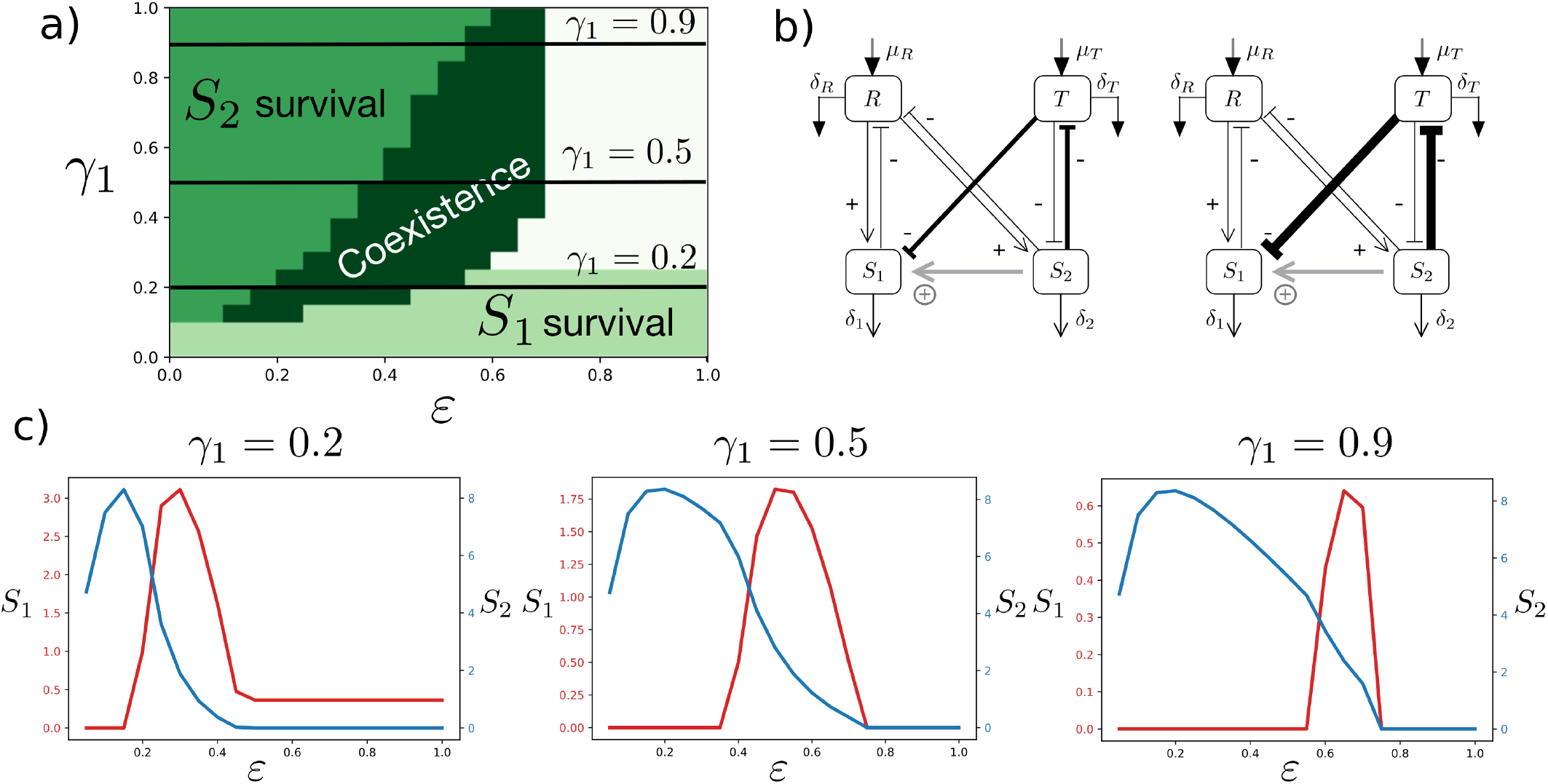
Positive correlation between the parameters regulating the indirect positive effect from species *S*_2_ to *S*_1_. a) Parameter space of the system as a function of *ε* and *γ*_1_ showing the different fixed points the system reaches. b) System scheme depicting the links of interest that follow a positive correlation. c) Density of species *S*_1_ and *S*_2_ (red and blue, respectively) for three conditions of *γ*_1_ (depicted also in a)) along the different values of *ε* that allows to observe the effect of the metabolic tradeoff parameter *ε* to the species’ fitness. The parameter space shown (*ε* vs. *γ*_1_) corresponds to the following constants: *μ*_*R*_ = *μ*_*T*_ = 0.5, *γ*_2_ = 0.1, *δ*_1_ = *δ*_2_ = 0.1, *ρ* = 0.3, *η* = 0.1, *δ*_*R*_ = *δ*_*T*_ = 0.1.

#### Toxic sensitivity asymmetry must hold to observe S_1_ and S_2_ coexistence

Although the model involves eleven parameters, just some have been found to be critical in order to understand its behaviour. The parameters determining the influx of *R* and *T* (*µ*_*R*_ and *µ*_*T*_, respectively) have been found significant to find regions of the parameter space where coexistence or survival of either of the species is possible (see Fig. 2). Importantly, from an evolutionary perspective, it is quite reasonable to conjecture that the resource and toxic waste levels could be of critical importance for such a system to emerge. Notice that, in general terms, the balance between *µ*_*R*_ and *µ*_*T*_ is crucial to determine the coexistence conditions of the species *S*_1_/*S*_2_. Similarly, the impact of the toxic on each species, weighted by *γ*_1_ and *γ*_2_, provides the other major influence.

When *γ*_1_*/γ*_2_ = 1, the impact of the toxic is symmetric: it reduces fitness in the same way for both strains. As shown below, the asymmetry between them will be the most relevant trait.

It is important to note that other parameters remain fixed and that a collapse of parameters can be made (see S-VII) without affecting the qualitative behaviour of the system. Resource *R* and toxic *T* degradation rates (*δ*_*R*_ and *δ*_*T*_) can be collapsed into *δ*, while cell mortality rates (*δ*_1_and *δ*_2_) can also be collapsed into a second parameter *δ*_*S*_. This notation will be used for the next derivations.

In Fig. 2 we summarise the main results of our analysis. Three particular conditions determine what we observe: [1]. *γ*_1_*/γ*_2_ ≤ 1, [2.] *γ*_1_*/γ*_2_ > 1 and [3]. *γ*_1_*/γ*_2_ ≫ 1. The asymmetry in these parameters (*γ*_1_ > *γ*_2_) is needed in order to find coexistence in the system. For *γ*_1_ ≤ *γ*_2_, coexistence is not possible even if other system’s parameters are changed. The linear boundary separating the presence of cells from its extinction can be easily derived in the latter case. Since *S*_2_ = 0, assuming that *R* and *T* achieve equilibrium much more rapidly than *S*_1_, and thus *dR/dt* ≈ 0 and *dT/dt* ≈ 0, we have *T* ≈ *µ*_*T*_ */δ* and also *R* ≈ *µ*_*R*_*/*(*δ*+*ηS*_1_) = *R*(*S*_1_) we can write a single-equation model for *S*_1_:

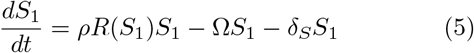

where we define Ω = *γ*_1_*µ*_*T*_ */δ* and *R*(*S*_1_) = *µ*_*R*_*/*(*δ* + *ηS*_1_), respectively. We can see that growth will occur only if *dS*_1_*/dt* > 0 or, in other words, if

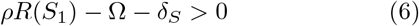

which leads, after some algebra, to the threshold condition between *µ*_*T*_ and *µ*_*R*_:

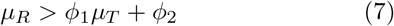

where *ϕ*_1,2_ are combinations of parameters, i. e. *ϕ*_1_ = *γ*_1_*/ρ* and *ϕ*_2_ = *δδ*_*S*_ */ρ* (see S-II SM for a more detailed analysis). This limit case thus predicts that the input of resources needs to be larger than the one associated to the toxic influx following a simple linear relation (consistently with the phase diagrams at left in Fig. 2). On the other hand, the larger the negative effect of *T* (as given by *γ*_1_) the more difficult for *S*_1_ to survive, as expected.

The critical threshold defined by the previous condition is also an estimate of the envelope of the parameter space where populations are expected to be found. The two other columns in the same figure indicate that if *γ*_1_ is larger than *γ*_2_ coexistence is achieved.

The original 4-equations model can be reduced, as we did before, to a two-dimensional system (showing the same behaviour as the 4D counterpart, see Fig. S2 and S3). Once again we assume that the dynamics of both *R* and *T* is faster than the ones associated to both *S*_1_ and *S*_2_ (see S-IV SM). A system of equations is then obtained, namely

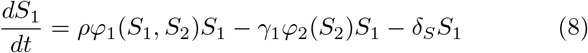

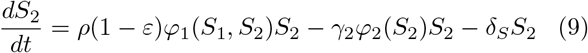

Here the two functions *φ*_*k*_ are given by:

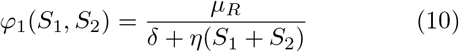

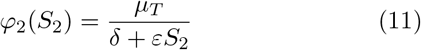

The two strains will be present provided that the two derivatives are *dS*_*k*_*/dt* > 0. This leads to two inequalities that need to be simultaneously satisfied, namely:

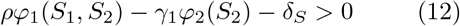

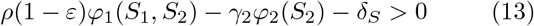

By dividing the two previous equations, it is easy to see that the following inequality is needed, namely we require:

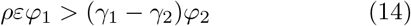

and, provided that *δ*_*S*_ is small enough compared with *S*_1,2_ the following condition for coexistence condition is found:

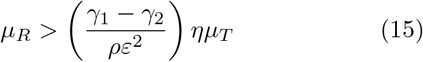

Which gives again a linear dependency that will hold only if asymmetry (with *γ*_1_ > *γ*_2_) is present. For the special case where *γ*_1_ ≫ *γ*_2_ we can actually neglect *γ*_2_ and write

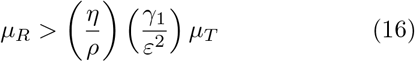

Another important feature that can be extracted from Fig. 2 is that the higher *γ*_1_ is with respect to *γ*_2_, the parameter space area with *S*_1_ survival vanishes and an area with *S*_2_ survival appears. The position of these particular areas is also of importance: *S*_1_ can survive on its own as long as *μ*_*T*_ remains low enough once its mortality rate due to toxic (*γ*_1_) is already slightly higher than *γ*_2_. On the other hand, the area with *S*_2_ survival appears on the right of the coexistence region: in this region, *μ*_*T*_ is considerably high and *S*_1_ has high sensitivity to *T,* thus only *S*_2_ survives. Finally, despite this is not specially relevant for the problem concerning us, not only the relative relation between *γ*_1_ and *γ*_2_ is relevant, but also their actual values. The higher *γ*_2_ is, the higher the difference with respect to *γ*_1_ must be in order to find coexistence of the species.

#### Positive correlation of indirect positive link parameter values

Along with the necessary coexistence condition *γ*_1_ > *γ*_2_, there is an extra condition to be fulfilled. We have found that the indirect positive relation between *S*_1_ and *S*_2_ (see Fig. 3) has a major role regulating the viability of the full system. This link has been depicted in Fig. 1c, formed by the toxic waste degradation by the species *S*_2_ (*−εTS*_2_) and the mortality induced to the species *S*_1_ by the toxic waste *T* (*−γ*_1_*TS*_1_).

In the inequality of eqn. 16 we have isolated as a factor in the right hand side of the equation the term *γ*_1_*/ε*^2^ that contains the inhibition *γ*_1_ of *T* on *S*_1_ as a proportional term along with *ε*, which acts here as an inverse square. Since *ε* is bounded between zero and one, its effects are stronger. This implies that relatively high values of epsilon must be counterbalanced by relatively lower (in comparison) values of *γ*_1_ in order that the inequality holds, keeping a nonlinear proportionality relationship. Therefore, the inequality 16 predicts in fact a positive correlation between these two parameters, which thus strongly influence the indirect positive effect.

The critical boundary separating coexistence from extinction in the (*γ*_1_, *ε*) space will scale as:

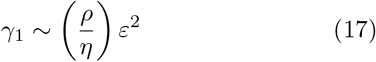

which of course only provides an (estimated) lower bound and predicts a nonlinear growth of *γ*_1_ with *ϵ*.

In Fig. 3a we can observe the actual relation between *ε* and *γ*_1_ that allows the coexistence of the species. A biological interpretation of this observation could be that, out of the region delimited by *ε* and *γ*_1_,the strength between toxic waste degradation by *S*_2_ and the mortality induced by *T* to *S*_1_ becomes too unbalanced, so that the indirect positive effect from *S*_2_ to *S*_1_ is too weak to actually sustain the survival of both species. We can also check in Fig. S11 how the positive relation between *ε* and *γ*_1_ prevails for various combinations of *μ*_*R*_ and *μ*_*T*_ values. Notice that if *ε/γ*_1_ ≫ 1, then *S*_2_ is unable to survive anymore due to its poor reproduction capacity (recall that the growth of *S*_2_ is regulated by *ρ*(1 - *ε*)*RS*_2_), and we find an area of the parameter space of *S*_1_ survival. On the other hand, if *ε/γ*_1_ ≪ 1, then the mortality induced by the toxic waste to *S*_1_ is too high for it to survive, and at the same time *S*_2_ has a much better reproduction capacity due to lower values of *ε*, so we find a region of the parameter space of *S*_2_ survival.

Fig. 3c provides more insights in what is actually occurring to the species densities across this parameter space. We can observe that either *S*_1_ and *S*_2_ have an optimal value of *ε* regarding its fitness. The value for *S*_2_ is located around 0.2, while the one for *S*_1_ is obviously dependent on *γ*_1_ but also, importantly, on the fitness of *S*_2_ associated to *ε*. Once *ε* keeps increasing and *S*_2_ degrades toxic with more efficiency and at the same time looses reproduction potential, *S*_1_ is able to survive and reach a peak of fitness at a concrete value of *ε*.

With this result we complete the explanation given above on the conditions for coexistence to be possible: coexistence is possible as long as *γ*_1_ > *γ*_2_ and, importantly, the values of *γ*_1_ and *ε* are confined into the region where the indirect positive interaction is strong enough to sustain it. Regarding the effect of other parameter values to the system’s behaviour, we have found that an asymmetry in cell mortality rates can lead to a subcritical Hopf bifurcation (see Fig. S12), but the qualitative behaviour of the system is maintained. Any other change in parameter values does not change the conclusions reached in this paper.

### B. Discrete Stochastic model

The mean field model is validated through the introduction of an spatial counterpart, where stochasticity is present, confirming the robustness of the species coexistence result under the scenario studied in this paper. In order to introduce noise in our model, we have implemented a discrete stochastic version of the mean field model in a 100×100 lattice. The rules are directly derived from the system of equations defined above, now considered to be discrete:

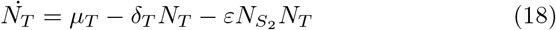

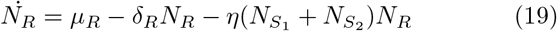

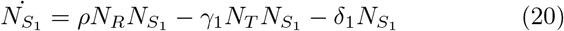

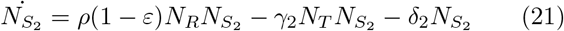

Each position (*i, j*) in the lattice contains discrete values of 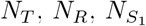 and 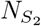 (*reactants*), and they will react locally with the reactants located in the same position (*i, j*). In this model there are up to three different types of reactions, depending on how many elements interact: [1]. influx of a certain element (just *N*_*R*_ and *N*_*T*_), [2]. *one-species* reaction proportional to a certain parameter value and [3]. *two-species* reaction when two elements interact also proportional to a particular rate. For a detailed description of the model rules see S-X SM and Fig. S13a.

The model includes random movement of 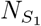 and 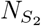 to its Moore neighbourhood of a given percentage of cell quantities (%_*move*_), as sketched in Fig. S13b, but does not consider discrete diffusion of *R* and *T* molecules.

The same parameters studied in the mean field model have been explored in the spatial counterpart, as can be observed in Fig. 4 (see Fig. S14 to check the actual levels of *S*_1_ and *S*_2_ under the conditions tested). The qualitative maintenance of the results obtained by the mean field model can be observed, as well as some changes in the steady state achieved with this stochastic counterpart. Namely, the state of species coexistence can be found in regions where it was precluded in the mean field model: either when γ_1_ < γ_2_ and when γ_1_*/*γ_2_ > 1, where before, under a critical value of *µ*_*T*_, only the survival of *S*_1_ was observed. This could be explained as a direct influence of the stochasticity now present in the simulations, due to each of the reactions aforementioned, the presence of the spatial dimension and the movement of the species *S*_1_ and *S*_2_ on it. In addition, the positive correlation between *ε* and *γ*_1_ is also confirmed in this stochastic version of the mean field model (see Fig. S15).

**FIG. 4:**
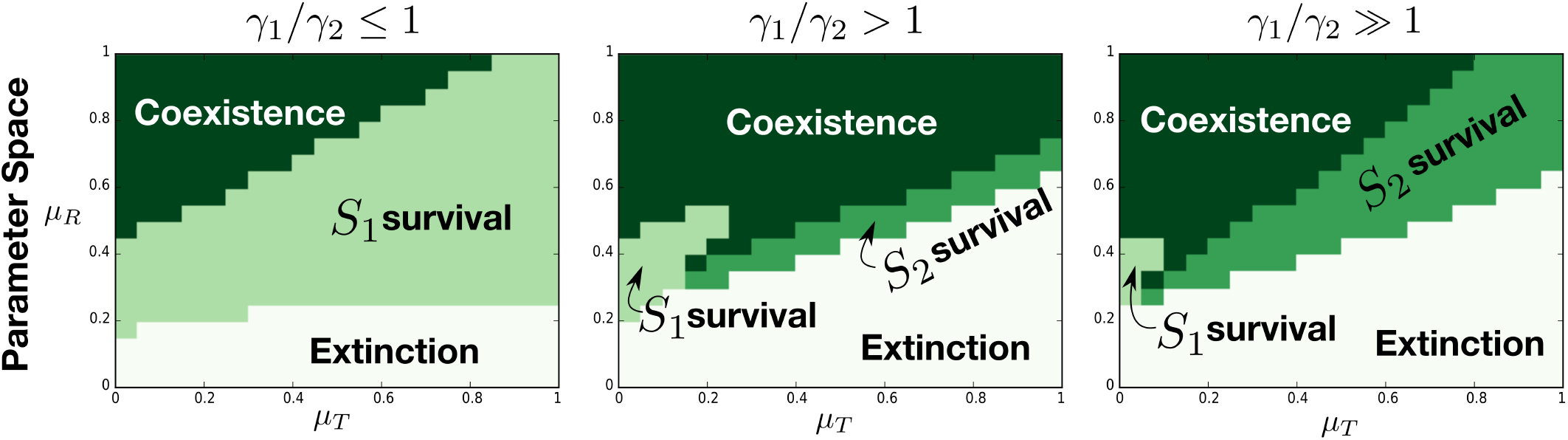
Parameter space for the discrete stochastic model under different conditions of *γ*_1_*/γ*_2_ (each column) as a function of *μ*_*T*_ and *μ*_*R*_. We observe how the results observed for the numerical simulations in Fig. 2 are qualitatively recovered, while some changes in the steady state are achieved with this stochastic counterpart in some regions of the parameter space. For *γ*_1_*/γ*_2_ ≤ 1, *γ*_1_ = *γ*_2_ = 0.1, for *γ*_1_*/γ*_2_ > 1, *γ*_1_ = 0.17 and *γ*_2_ = 0.1 and for *γ*_1_*/γ*_2_ ≫ 1, *γ*_1_ = 0.3 and *γ*_2_ = 0.1

The levels of *R, T, S*_1_ and *S*_2_ rapidly reach a stable mean value, around which each value of the four system dimensions in every position of the lattice (*i, j*) oscillates (see Fig. S16 for a sample of the temporal evolution of the mean values ± SD of each dimension).

## IV. DISCUSSION

In this paper we explore the persistence of stable heterogeneous cell assemblies under the presence of an environmental challenge as defined by a toxic waste. To this goal we consider a simple mathematical model (and its stochastic counterpart) where two cell populations exploit available resources while being constrained by the lethal effects of the toxic. This situation can be relevant in different contexts, from evolutionary dynamics of multicellularity to the maintenance of diverse populations under heterogeneous physiological conditions (as it occurs with biofilms[27]). Given that the production of metabolites inevitably leads to some kind of inhibitory signals, it is relevant to consider how a diverse population can persist.

In our system, one population is capable of removing the toxic waste at the expense of a reduced reproductive potential under a metabolic tradeoff. This scenario is related to early studies of competition in microbial communities [14–16] where a related problem was addressed in the context of antibiotic resistance. But in those models the toxic inhibitory species was considered to be resistent to it while no metabolic tradeoffs where present. These are important features of a recent theoretical work where the emergence of complex, spatially distributed “proto-organisms” (PRO) exhibiting division of labor and higher-scale features occurs[12], now taken into account in the presented mean field model. Beyond the differences with the original PRO model, we take the most fundamental features of it in order to understand the possible key features predating the observed complex PRO.

Our analysis provides robust predictions regarding the conditions for stability. As a consequence of the interaction network resulting from our set of rules, well-defined domains of persistent mixed species is found. A theoretical characterisation is provided for the continuous, well-mixed model, while a discrete spatial model is also analysed to test the robustness of the observed phases. Under this scenario, we also show that an asymmetry between their sensitivities to toxic must exist. In particular the sensitivity to toxic for the toxic-degrading species *S*_2_ must be lower than the *wild type* species *S*_1_. In an evolutionary context, this implies that the mutation into a new species capable of degrading a certain toxic waste should display a lower sensitivity to it (compared to the *wild type* species) in order to allow coexistence. Importantly, this conclusion holds when stochasticity is at play.

On the other hand, we show how the indirect positive relation from the toxic degrading species to the *wild type* species (which could be classified as a commensalist relationship) is critical for the existence of cell heterogeneity. A positive correlation between the parameters regulating the strength of the two links conforming the indirect positive relation must hold in order to observe species coexistence. Again, the spatial stochastic model confirms the robustness of this result. It is plausible that this kind of relationship between two related species through a toxic species could be potentially selected in an evolutionary context where an increase in complexity in the form of cell heterogeneity is selected for. Moreover, the presence of a metabolic tradeoff for the toxic-degrading species could further constrains the viability of both species in this evolutionary context, as it has been shown in the paper.

In conclusion, we provide a detailed analysis of a new scenario of toxic-mediated cell heterogeneity with potential for synthetic purposes. With the aim of studying the origins of multicellularity, a theoretical framework has been recently proposed[28], where the use of synthetic multicellularity as a tool to explore the possible in terms of *Ecology, Evolution* and *Development* was presented. In this paper we use the *ecological* dimension to study a potential system that could predate the further evolution of cells into complex multicellular structures. Our results could be of potential use for synthetic biologists aiming to design a system where two cells coexist having a commensalist relationship. This basic system could then be synthetically evolved (studying the proposed *Evolution* dimension) to explore possible paths to generate stable multicellular aggregates probably underlying mutualistic relationships. These potential (both theoretical and experimental) extensions will require further investigation.

## Data Accessibility

The results have been generated through the widely known Runge Kutta numerical method (see SXI SM) while the Discrete Stochastic model can be easily reproduced with the information given in the paper (section III-B in main text and S10 in SM).

## Research ethics

This statement does not apply to this manuscript.

## Animal ethics

This statement does not apply to this manuscript.

## Permission to carry out fieldwork

This statement does not apply to this manuscript.

## Authors’ contributions

RS conceived the idea and design of this work, analysed and interpreted results and wrote the article. AOV helped in the design of the work, developed the code, performed the experiments, analysed and interpreted results and wrote the article.

## Acknowledgments

The authors thank the members of the Complex Systems Lab for useful discussions. Special thanks to Rosa Martínez-Corral and Jordi Piñero for our discussions and their valuable suggestions and to G. Taro for her inspiring ideas.

